# Plant architecture is correlated with variation in cannabinoid concentration and biomass production in *Cannabis sativa*

**DOI:** 10.1101/2022.12.02.518916

**Authors:** George M. Stack, Craig H. Carlson, Jacob A. Toth, Glenn Philippe, Jamie L. Crawford, Julie L. Hansen, Donald R. Viands, Jocelyn K. C. Rose, Lawrence B. Smart

## Abstract

*Cannabis sativa* is cultivated for multiple uses including the production of cannabinoids. In developing improved production systems for high-cannabinoid cultivars, scientists and cultivators must consider the optimization of complex and interacting sets of morphological, phenological, and biochemical traits, which have historically been shaped by natural and anthropogenic selection. Determining factors that modulate cannabinoid variation within and among genotypes is fundamental to developing efficient production systems and understanding the ecological significance of cannabinoids. Thirty-two high-cannabinoid hemp cultivars were characterized for traits including flowering date and shoot-tip cannabinoid concentration.

Additionally, a set of plant architecture traits, as well as wet, dry, and stripped inflorescence biomass were measured at harvest. One plant per plot was partitioned post-harvest to quantify intra-plant variation in inflorescence biomass production and cannabinoid concentration. Some cultivars showed intra-plant variation in cannabinoid concentration, while many had a consistent concentration regardless of canopy position. There was both intra- and inter-cultivar variation in architecture that correlated with intra-plant distribution of inflorescence biomass, and concentration of cannabinoids sampled from various positions within a plant. These relationships among morphological and biochemical traits will inform future decisions by cultivators, regulators, and plant breeders as well as our broader understanding of intra-plant variation of specialized biochemicals.

**Highlight:** In-season hemp plant architecture measurements can predict post-harvest traits related to the distribution of biomass and concentration of cannabinoids.

## 1 Introduction

Plant secondary metabolism generates diverse classes of chemical compounds that have numerous ecological and anthropic functions. Many of these compounds mediate interactions between plants and their environment (Bennett and Wallsgrove, 1994; Wink, 2008; Kessler and Kalske, 2018), and more recent studies have demonstrated their complex multifunctionality and utility in primary metabolic processes (Erb and Kliebenstein, 2020). Ecological theory predicts ontogenetic and tissue-specific variation in the concentration of plant secondary metabolites, primarily in the context of herbivore resistance (McKey, 1974; Pavia *et al*., 2002; van Dam, 2009; Meldau *et al*., 2012; Schuman and Baldwin, 2016; Barton and Boege, 2017). There are many examples of ecologically relevant intra-organismal variation in secondary metabolite concentration across diverse taxa, including *Gossypium* (Anderson and Agrell, 2005), *Brassica* (Gutbrodt *et al*., 2012), *Acer* (Mason *et al*., 2019), *Populus* (Mason *et al*., 2019), *Caulerpa* (Meyer and Paul, 1992), *Dictyota* (Cronin and Hay, 1996), and *Oceanapia* (Schupp *et al*., 1999). The well-studied cases of biochemically-mediated plant resistance to herbivores provide insight into the evolutionary drivers responsible for heritable intra-plant variation in secondary metabolites.

*Cannabis sativa* synthesizes and stores cannabinoids, a class of secondary metabolites, in glandular trichomes that are most densely produced on female inflorescences (Dayanandan and Kaufman, 1976; Turner *et al*., 1978; Livingston *et al*., 2020). The adaptive value of cannabinoid synthesis prior to human cultivation is not known, but theories include protection from herbivores, pathogens, or ultraviolet radiation (Pate, 1983; Lydon *et al*., 1987; Gorelick and Bernstein, 2017; Tanney *et al*., 2021). In any of these cases, one might predict that intra-plant variation in cannabinoid production, with greater concentrations of cannabinoids in tissues with a greater fitness value, would be evolutionarily advantageous, as cannabinoids are costly to synthesize. Consistent with this prediction, Richins et al. (2018) and Bernstein et al. (2019) reported significantly greater concentrations of cannabinoids in inflorescences than in vegetative leaves; however, this may be, in part, a result of strong human-driven selection for cannabinoidrich inflorescences.

Beyond the hypothesized adaptive value, cannabinoid concentration is among the most important factors considered by commercial producers and processors of high-cannabinoid *C. sativa*. The two major harvested products of high-cannabinoid *C. sativa* are bulk inflorescence biomass for extraction and trimmed smokable flowers for direct consumption. For producers of high-cannabinoid hemp, the concentrations of cannabinoids, specifically Δ^9^□tetrahydrocannabinol (THC) and tetrahydrocannabinolic acid (THCA), are of particular importance in the context of regulatory compliance (USDA-AMS, 2021). In most jurisdictions, there is a legislated threshold of total potential THC concentration that distinguishes hemp from marijuana, and if exceeded, growers risk losing both their crop and license to grow hemp.

The samples for regulatory testing for THC compliance and ‘potency’ testing to determine the value of biomass or smokable flower are collected at various time points of the crop production and post-harvest processing pipeline. Regulatory tissue samples in hemp production are typically taken from the top of an inflorescence several weeks prior to harvest, but protocols vary by jurisdiction. Biomass is often sampled after drying and stripping the floral material from stems and branches. However, this process is often imprecise, and the low-cannabinoid content of residual stem and leaf biomass can dilute the concentration in a bulk sample. Smokable flower samples are often tested after the inflorescence has been trimmed of leaf tissue, leaving predominately trichome-rich perigonal bracts.

Previous studies have demonstrated that some *C. sativa* genotypes exhibit intra-plant variation in cannabinoid concentration. Richins et al. (2018) and Bernstein et al. (2019) both quantified cannabinoids from various regions of the canopy, finding generally greater concentrations of cannabinoids in samples from the upper canopy than from the lower canopy. These findings align with anecdotal evidence from high-cannabinoid *C. sativa* producers who consider the primary apical inflorescence as being the most cannabinoid-rich. Additionally, many of these producers impose labor-intensive plant architecture manipulations through pruning with the goal of increasing inflorescence biomass production and cannabinoid concentration, while mitigating biotic pressures.

The interaction between canopy architecture and canopy position on cannabinoid concentration has recently been investigated by measuring the effects of various pruning treatments. Crispim Massuela et al. (2022) sampled inflorescences from three canopy positions (upper, middle, and lower) of plants that were un-pruned, had apical meristems cut back, or had the lower canopy branches removed. They found that only inflorescence position, not pruning technique or its interaction with inflorescence position, had a significant effect on total cannabidiol (CBD) concentration. Danziger and Bernstein (2021) imposed eight different pruning treatments consisting of various combinations of leaf and branch removal from various parts of the canopy, and computed a ‘plant uniformity score’ to quantify deviation from the mean concentration of cannabinoids among inflorescences. They found that pruning treatments can increase the computed plant uniformity score through small cannabinoid concentration increases in inflorescences that were shaded in the un-pruned control. Pruning either decreased or did not change the concentration of total THC, the predominate cannabinoid, in the primary apical inflorescence. These studies provide important insight into the effect of canopy architecture on intra-plant variation in cannabinoid production. However, additional related studies of a comparatively more diverse set of *C. sativa* germplasm are needed to validate these findings, as well as account for potential genotype-by-architecture interactions.

The primary objectives of this study were to (1) characterize a set of high-CBD hemp cultivars for in-season morphological and biochemical traits, (2) characterize the partitioning of stripped inflorescence biomass throughout the plant canopy, (3) quantify intra-plant variation in cannabinoid concentration, and (4) evaluate the relationships between morphological and biochemical traits to inform ecological models, as well as future decisions by hemp cultivators, regulators, and breeders.

## 2 Materials and Methods

### 2.1 Plant material and propagation

Thirty-two high-CBD hemp cultivars (Table 1) from nine commercial sources and the Cornell hemp breeding program were established in a peat-based soilless media (Lambert LM-111) in the second week of May 2020. In accordance with techniques employed by commercial cultivators, plants were propagated from dioecious (male and female) seed, ‘feminized’ (all female) seed, or via two-node vegetative cuttings. Dioecious cultivars were screened at the seedling stage with the *C. sativa* Y chromosome-specific molecular marker, CSP-1 (Toth *et al*., 2020), in order to select exclusively females for transplant to the field. Cuttings were rooted using Clonex^®^ rooting hormone (Hydrodynamics International, Lansing, MI). Seedlings and cuttings were maintained in the greenhouse at 18 h light:6 h dark until transplant in the first week of June.

**Table 1.**
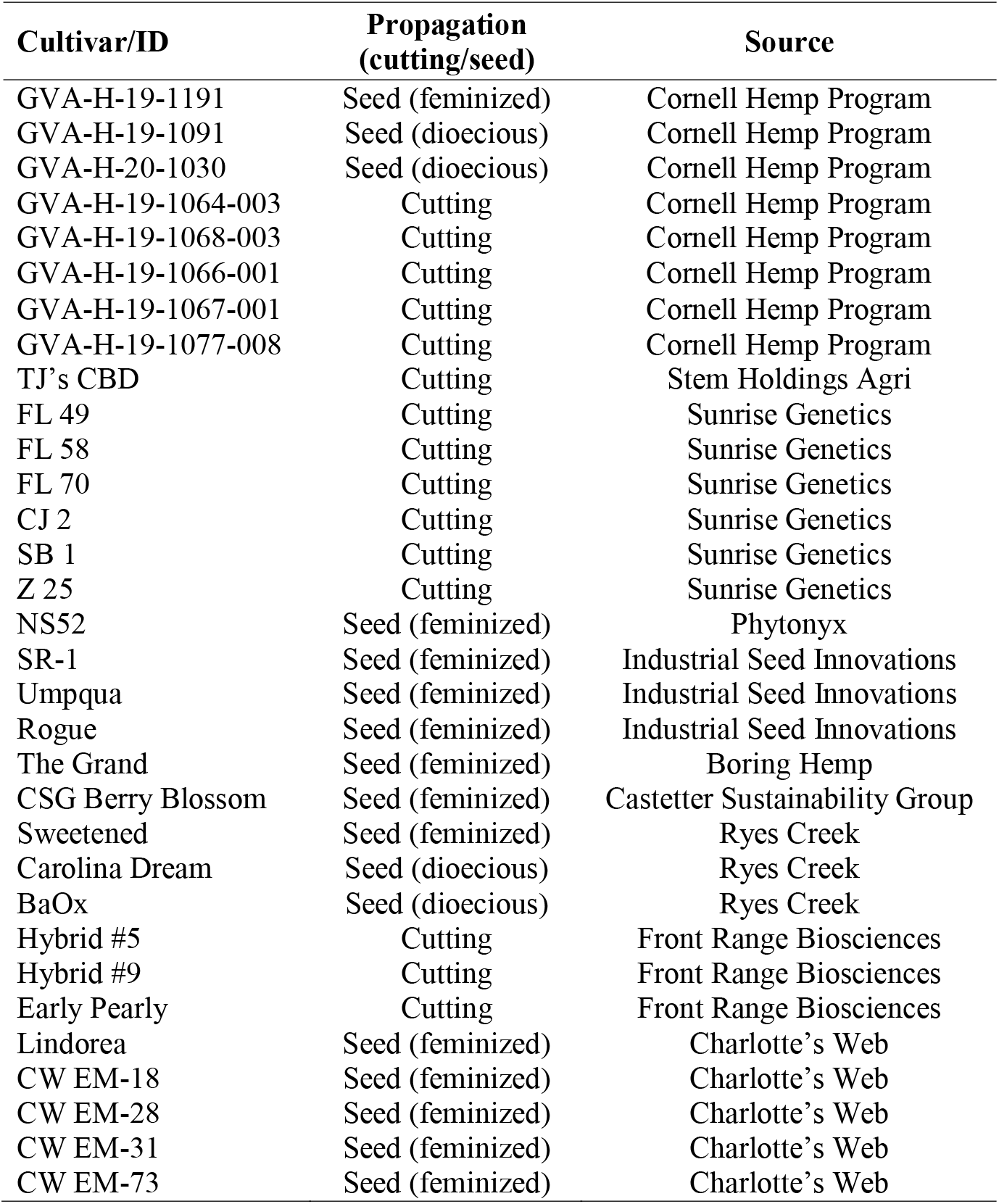
Sources of 32 high-CBD cultivars included in field trials in Geneva and Ithaca. Cultivars were started from seed (dioecious or feminized) or vegetative cuttings. Cultivars were generously contributed by the companies indicated.

### 2.2 Field preparation and maintenance

Trials were planted at two Cornell University field sites: Geneva, NY (McCarthy Farm: 42.9, -77.0) and Ithaca, NY (Bluegrass Lane Turf and Ornamental Farm: 42.5, -76.5). Weather station data can be found at http://newa.cornell.edu (Last accessed on 11/22/2022). Each site was cultivated and raised beds formed with drip irrigation and black plastic mulch prepared every 1.83 m on center. Fertilizer (19-19-19, Phelps Supply Inc., Phelps, NY) equivalent to 95.3 kg N ha^-1^, was spread under the plastic mulch in Geneva and was broadcast pre-planting in Ithaca. Landscape fabric was deployed to control weeds in alleys between rows.

Each cultivar was planted in five-plant plots in a replicated, randomized complete block design with four replicate blocks at each site. Seedlings and rooted cuttings were transplanted into raised beds on June 11, 2020 (Geneva, NY) and June 12, 2020 (Ithaca, NY). Plants were spaced 1.23 m apart within rows. After transplanting, the plots were irrigated using in-bed drip irrigation as needed throughout the season to maintain optimal soil moisture (>0.27 m^3^ m^-3^). HOBOnet RXW-SMD-10HS soil moisture sensors (Onset, Bourne, MA) were installed at a depth of 10 cm and used to assess when irrigation was necessary. Fertilizer (Jack’s 12-4-16 Hydro FeED RO, 11.3 kg per application) was included in the irrigation on two occasions in early and late July.

### 2.3 In-season measurements

The heights of the middle three plants of each five-plant plot were measured weekly after transplant until there was no change in height for two consecutive weeks. All plants were surveyed weekly to determine the onset of flowering following the protocol described by Carlson et al. (2021), where ‘terminal flowering’ describes distinct clusters of pistillate flowers observed at shoot apices (Supplementary Fig. S1). Plants that produced staminate flowers were promptly removed from the field to maintain unpollinated female inflorescences.

For cannabinoid quantification, regulatory-style shoot-tip samples of the 32 cultivars were collected from a single plant in every plot starting one week after terminal flowering and again at three and five weeks after terminal flowering. In accordance with regulatory standard in New York State at the time of the trial, the top 10 cm of the apical inflorescence was sampled for the time series. The first plant in the plot was sampled for the 1-week timepoint, the fifth for the 3-week timepoint, and the second for the 5-week timepoint. Shoot tip samples were dried in a climate-controlled room (∼30% RH and below 33**°**C) and then milled to a fine powder in a Ninja® Pro blender (SharkNinja, Needham, MA). Milled samples were stored at 4°C prior to high pressure liquid chromatography (HPLC) analysis following the methods described by Stack et al. (2021) (Supplementary Dataset S2). The following cannabinoids were quantified for each sample: tetrahydrocannabinolic acid (THCA), Δ^9^□tetrahydrocannabinol (THC), cannabidiolic acid (CBDA), cannabidiol (CBD), cannabichromenic acid (CBCA), cannabichromene (CBC), cannabigerolic acid (CBGA), cannabigerol (CBG), cannabinol (CBN), tetrahydrocannabivarin (THCV), tetrahydrocannabivarinic acid (THCVA), cannabidivarin (CBDV), cannabidivarinic acid (CBDVA), cannabicyclol (CBL), cannabicyclolic acid (CBLA), and Δ^8^□tetrahydrocannabinol (Δ^8^-THC). To control for variation in decarboxylation of acid-form cannabinoids and cultivar-dependent variation in relative proportions of different cannabinoids, the statistical analysis was performed on the sum of all total potential cannabinoid percentages. See Table 3 in Stack et al. (2021) for formulas used to calculate total potential cannabinoids.

### 2.4 Time-of-harvest and post-harvest measurements

At harvest, 5 weeks after terminal flowering, canopy architecture traits of the third plant in each plot were measured following the method described by Carlson et al. (2021), including height, maximum canopy diameter (MCD), and the height at the maximum canopy diameter (MCDH). Following the measurements, the stems were cut at soil level and the total wet biomass of each plant in a plot was measured. The third plant in each five-plant plot was partitioned into five sections (S1 through S5) based on the length of the main stem (Fig. 1). Sections were subsequently air-dried in a greenhouse with industrial fans.

**Figure 1.**
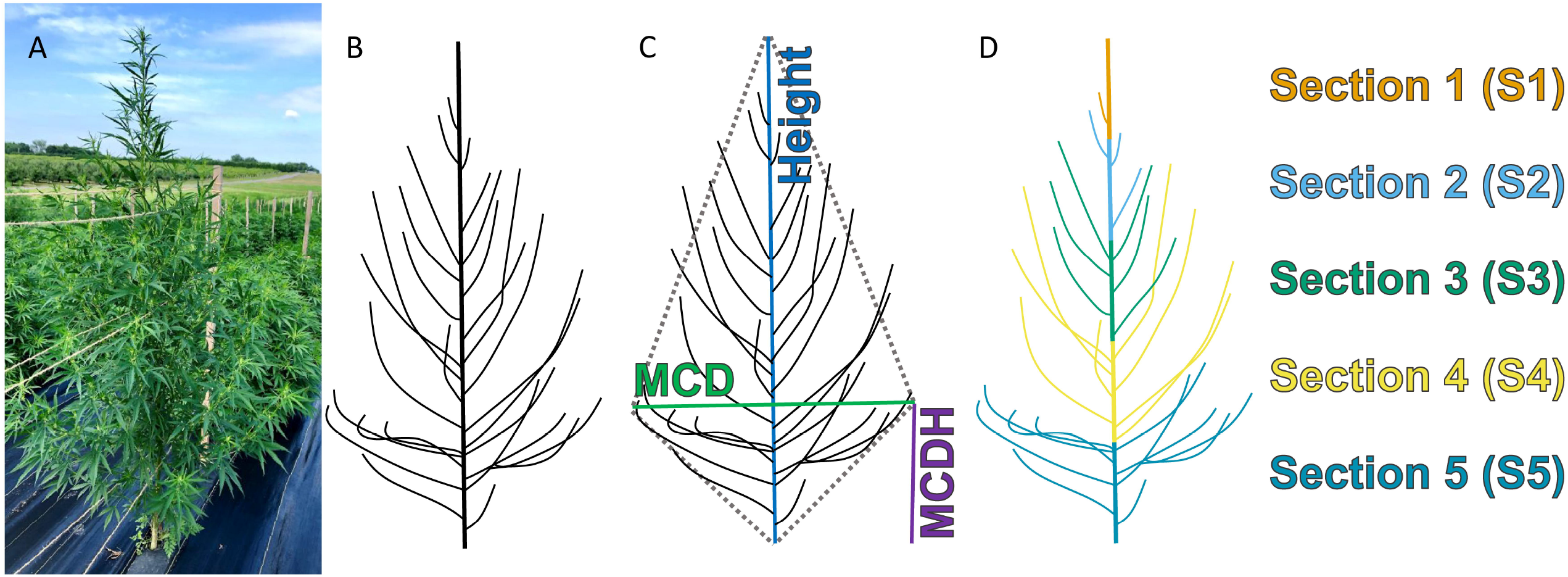
Schematic of (A) a hemp plant in the field, (B) the branching structure of the hemp plant, (C) the canopy-level plant architecture measurements taken prior to harvest, and (D) the sectioning scheme used to partition the plant into five sections.

After drying, basal stem diameter, total dry biomass – including stems, leaves, and inflorescence material – and dry stripped inflorescence biomass were measured for each section. The full list of traits measured with relevant formulas can be found in Supplementary Table S1. Cannabinoids were quantified by HPLC following the protocol above using a subsample of the homogenized stripped biomass for each section (Supplementary Dataset S3).

### 2.5 Statistical analysis

All statistical analyses and preparation of graphs were conducted in the open-source statistical computing platform R version 4.1.3 (R Core Team, 2022). Simple linear regression was used to determine cultivar-wise correlation and regression coefficients among cannabinoid sampling timepoints and positions. The cultivar-level mean proportions of dry biomass in each section were used to conduct a *k*-means clustering analysis following the Hartigan and Wong algorithm (Hartigan and Wong, 1979). The number of clusters (*n*=4) was determined by the elbow method (Supplementary Fig. S2). Simple linear regression was used at both the cultivar- and plot-level to determine correlations between the following values: the proportion of dry biomass in S5 and stripped inflorescence biomass per unit area, the MCD to height ratio and the proportion of dry biomass in S5, the MCD to height ratio and the log of the ratio between stripped inflorescence biomass produced in S4 and S5. Second-degree polynomial regression was used to model cultivar- and plot-level relationships between bicone volume and harvest index.

A mixed-effects model with biomass section and cultivar as fixed effects, and plot as a random effect, was used to determine whether a cultivar or biomass section had a significant effect on total cannabinoid concentration. A series of mixed-effects model F-tests were used to determine whether biomass section had a significant effect on total cannabinoid concentration for each cultivar. For all mixed-effects models a Satterthwaite approximation was used to estimate the effective degrees of freedom. When the effect of section was deemed significant after correcting for multiple testing using a Bonferroni correction (α = 0.0016), a post-hoc Tukey’s HSD analysis was used to test pairwise differences between sections.

To follow-up on the results of previous studies that reported intra-plant variation in cannabinoid concentration, cultivar-level cannabinoid concentration was regressed on plant section to calculate a slope corresponding to the change in cannabinoid concentration between sections from the top and moving down each section. Based on the results of Danziger and Bernstein (2021), canopy area, dry canopy density, and an interaction term were used to model the change in cannabinoid content by section at the cultivar level. The dry canopy density was log transformed to normalize the residuals, and non-significant terms were removed from the model before reporting model statistics.

Stepwise regression was performed with the function ‘step’ (direction=‘both’) using all biomass and plant architecture variables found in Supplementary Dataset S1 to model change in cannabinoid concentration by section at the cultivar-level. The package relaimpo (Groemping, 2007) was used to order predictors and calculate LMG indices to partition additive properties of *R*^2^. A mixed-linear model was used to predict change in cannabinoid concentration by section using the log of dry canopy density, canopy area, and an interaction term as predictors.

## 3 Results

### 3.1 Cannabinoid concentrations in regulatory samples are well correlated with concentrations in end-of-season biomass

Total potential cannabinoid concentration was positively correlated among all in-field sampling timepoints and postharvest biomass samples (Fig. 2, *p* < 0.05). The correlations between regulatory-style shoot samples and stripped biomass samples were strong, with *R*^2^ values ranging from 0.64 to 0.85. The concentration in end-of-season stripped bulk biomass was more strongly correlated with the concentration in samples from the biomass sections than the concentration in regulatory-style shoot tip samples. Samples from biomass sections generally had a greater correlation coefficient with those from adjacent sections, rather than with those from more distal sections. Samples from the bulk biomass had a greater correlation coefficient with lower sections of the canopy than with samples from upper sections.

**Figure 2.**
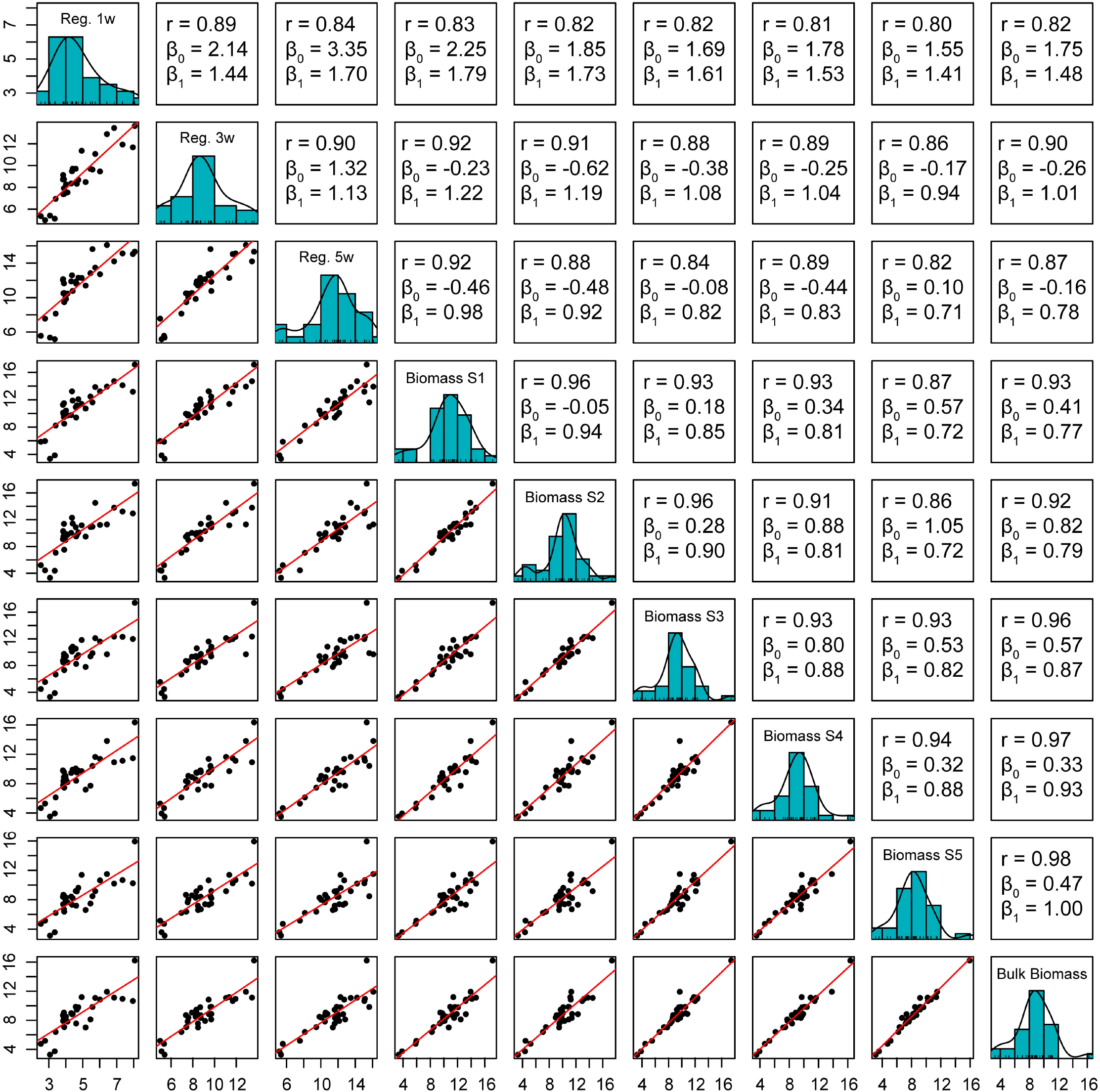
Cultivar-level pairwise correlations among total potential cannabinoid concentrations from samples over the course of floral maturation and harvest. The main diagonal is the distribution of data for the nine sampling points, from left to right: regulatory-style sample 1 week, 3 weeks, and 5 weeks after terminal flowering, post-harvest stripped biomass samples from plants partitioned into five sections where S1 is branches derived from the top 5^th^ of the main stem, S2 those from the second to top, etc., and bulk inflorescence biomass from the entire plant. Correlation and regression coefficients correspond to the regressions plotted in panels mirrored by the main diagonal.

### 3.2 Biomass distribution among sections is cultivar-dependent

The cultivars evaluated produced a wide range of stripped inflorescence biomass on a per-plant basis (Fig. 3A). Further, the proportion of total stripped biomass produced in each section of the canopy varied by cultivar (Fig. 3B). On one extreme, over 75% of the total stripped biomass produced by GVA-H-19-1077-008 was in S5, the section closest to the base of the plant. In contrast, less than 30% of the total biomass produced by ‘Umpqua’ was in S5. *K*-means clustering analysis of the proportions of dry stripped biomass in each section separated the cultivars into four clusters (Fig. 3B). The clusters corresponded almost perfectly with the variation in the proportion of stripped biomass in S5.

**Figure 3.**
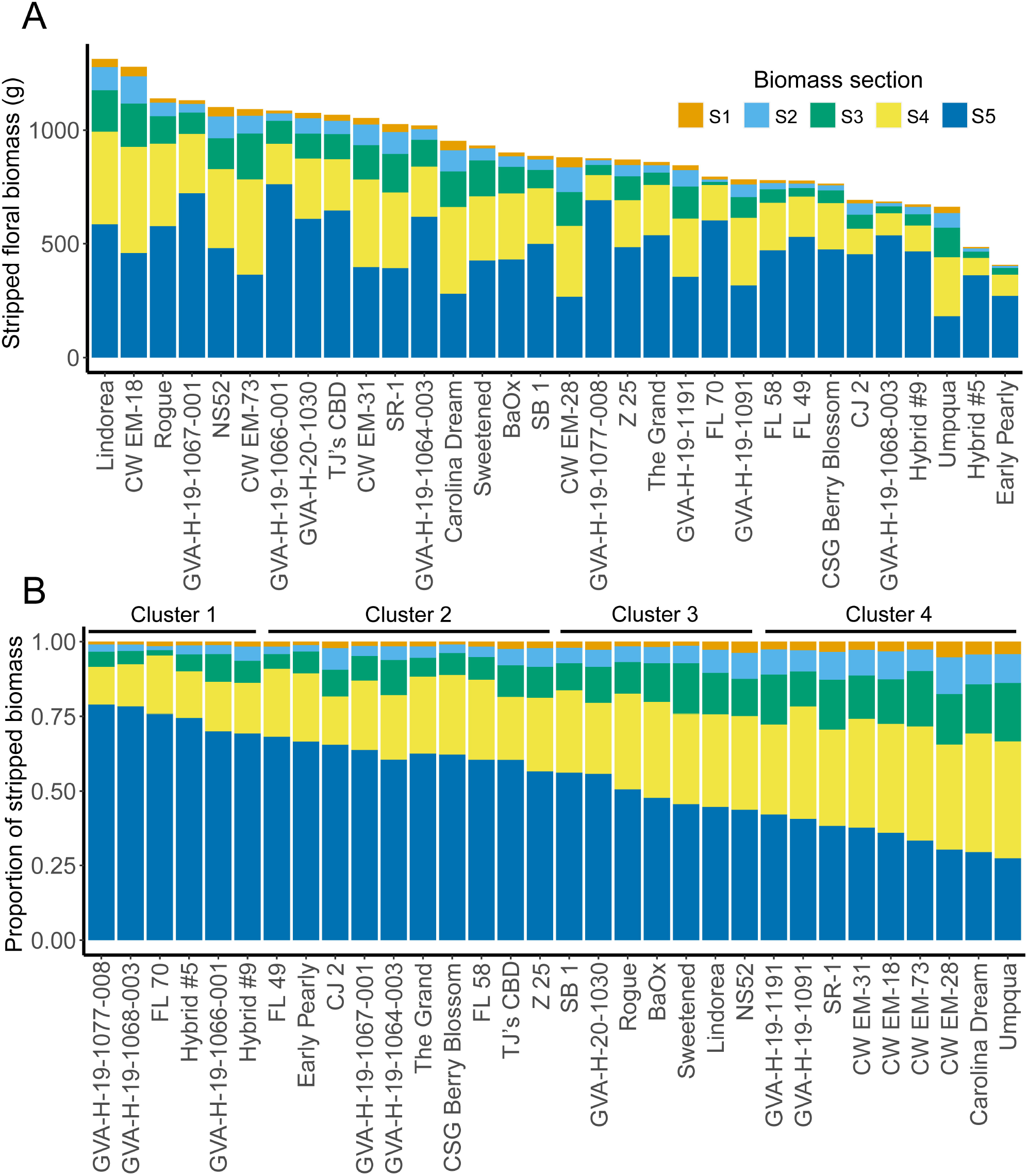
Bar plots of stripped biomass produced by 32 high-CBD hemp cultivars partitioned into five sections where S1 is branches derived from the top 5^th^ of the main stem, S2 those from the second to top, etc. Colors indicate the mean stripped biomass per plant (A) and the proportion of stripped biomass (B) from each of the five sections. Clusters in (B) are cultivar-level *k*-means clusters based on the proportions of dry biomass produced in each section.

### 3.3 Variation in plant architecture is correlated with biomass distribution

There was substantial variation in plant architecture among and within the 32 cultivars evaluated with plant height and MCD ranging from 61 cm to 249 cm and 55 cm to 265 cm, respectively (Fig. 4, Supplementary Dataset S1). The ratio of the MCD to height was positively correlated with the proportion of dry biomass in S5 at the level of cultivar (*R*^2^ = 0.47, *F*(1,30) = 26.1, *p* < 0.001) and plot (*R*^2^ = 0.29, *F*(1,228) = 91.35, *p* < 0.001) (Fig. 5), and negatively correlated with the log of the ratio between stripped biomass in S4 and S5 at the level of cultivar (*R*^2^ = 0.44, *F*(1,30) = 23.37, *p* < 0.001) and plot (*R*^2^ = 0.20, *F*(1,228) = 55.28, *p* < 0.001) (Fig. 5). There was a significant relationship between bicone volume and harvest index at the level of cultivar (*R*^2^ = 0.57, *F*(2,29) = 55.28, *p* < 0.001) and plot (*R*^2^ = 0.39, *F*(2,227) = 73.11, *p* < 0.001) (Fig. 4). The proportion of dry biomass in S5 was negatively correlated with the log of the amount of stripped inflorescence biomass per unit area at the level of cultivar (*R*^2^ = 0.33, *F*(1,30) = 14.56, *p* < 0.001) and plot (*R*^2^ = 0.09, *F*(1,228) = 23.81, *p* < 0.001) (Fig 5).

**Figure 4.**
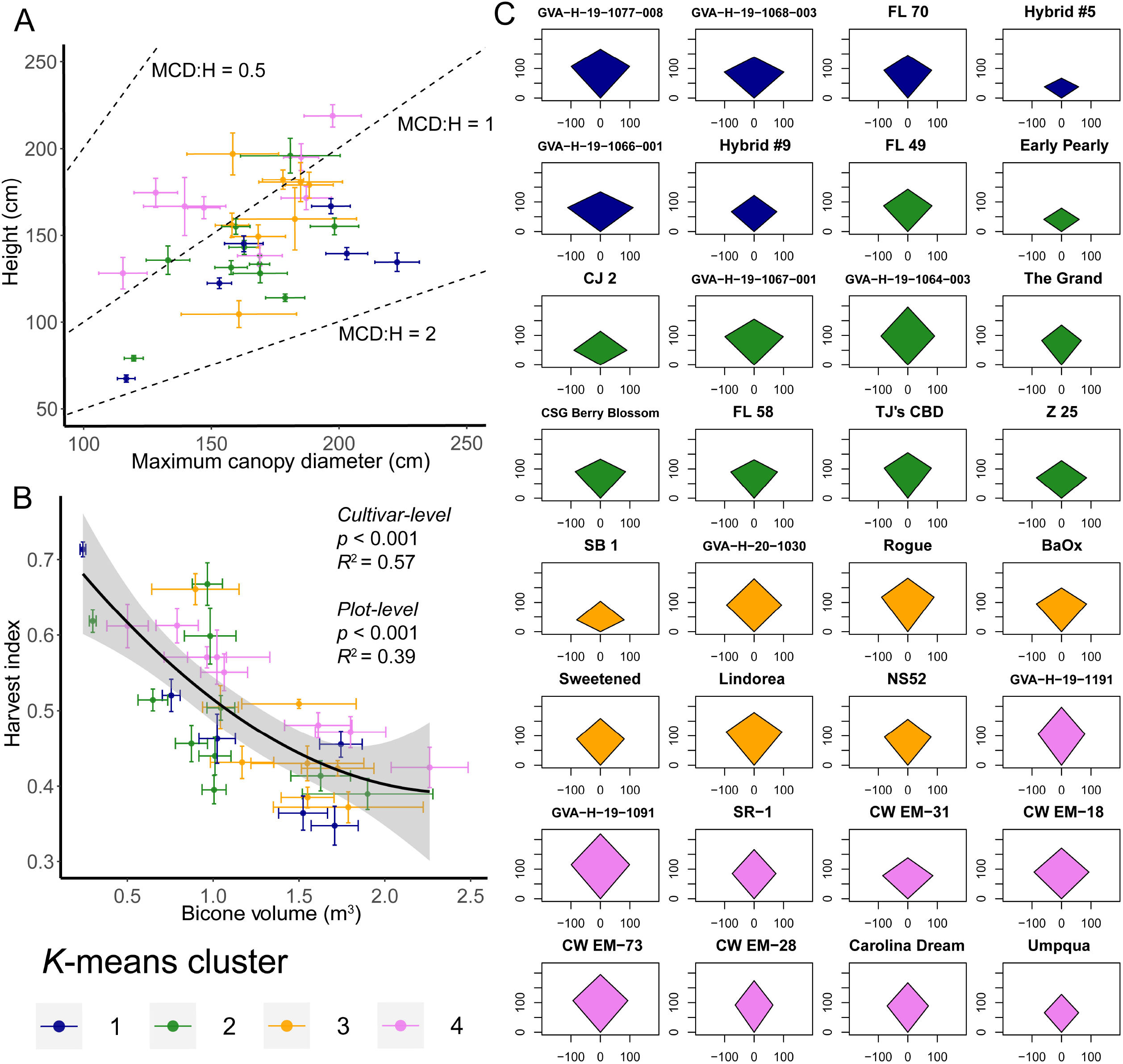
Variation in plant architecture within and among high-cannabinoid hemp cultivars. (A) plot of maximum canopy diameter (MCD) versus height where points are cultivar means and error bars are the standard error of the mean. Dashed lines indicate MCD to height ratios of 0.5, 1, and 2. (B) Second-degree polynomial regression of harvest index on bicone volume where points are cultivar means and error bars are the standard error of the mean. (C) Average kite models by cultivar contrasted from MCD, height, and the height of the maximum canopy diameter (MCDH). Colors in all panels indicate the *k*-means cluster to which that a cultivar belongs based on the proportions of dry biomass in each biomass section.

**Figure 5.**
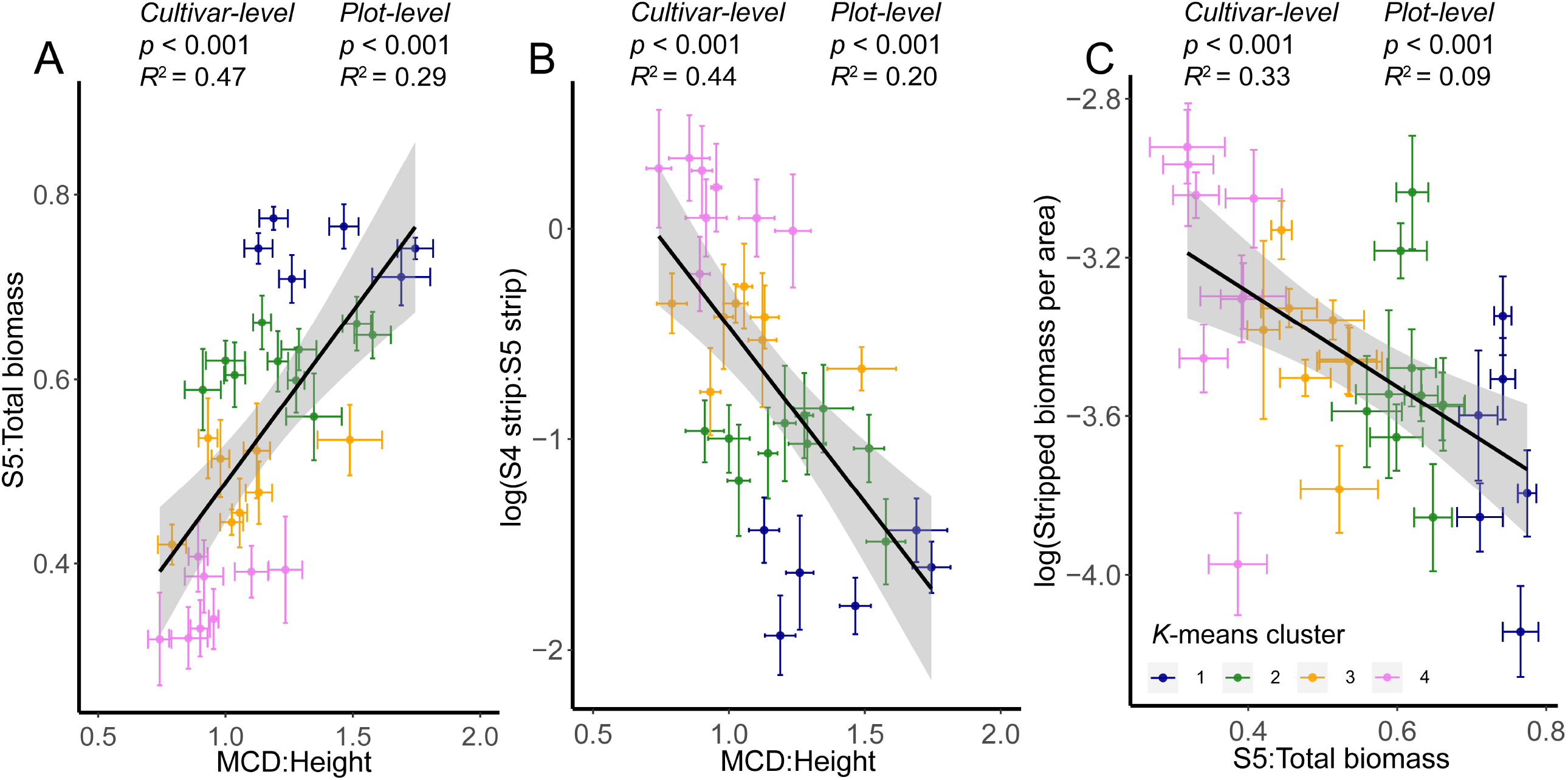
Correlations between (A) MCD:Height ratio and proportion of dry biomass in S5, (B) MCD:Height ratio and log of the ratio of stripped biomass produced in S4 and S5, and (C) proportion of dry biomass in S5 and log of stripped biomass per unit area. Each point and set of error bars, depicting the standard error of the mean, represents one cultivar. Colors indicate

### 3.4 Intra-plant variation in cannabinoid concentration is cultivar-dependent

When modeling the concentration of total potential cannabinoids sampled from the biomass sections using a mixed-effects model, there were significant main effects of cultivar (*F*(31,502.12) = 16.23, *p* < 0.001) and biomass section (*F*(1,988.23) = 434.14, *p* < 0.001), and a significant interaction between cultivar and biomass section (*F*(31,988.22) = 5.24, *p* < 0.001). After correcting for multiple testing for the 32 mixed-effects model F-tests (α = 0.0016), 17 of the cultivars had statistically significant variation in total cannabinoid concentration among plant sections (Fig. 6). For those cultivars with significant variation among sections, the cannabinoid concentration in S1 was always significantly greater than the concentration in S5.

**Figure 6.**
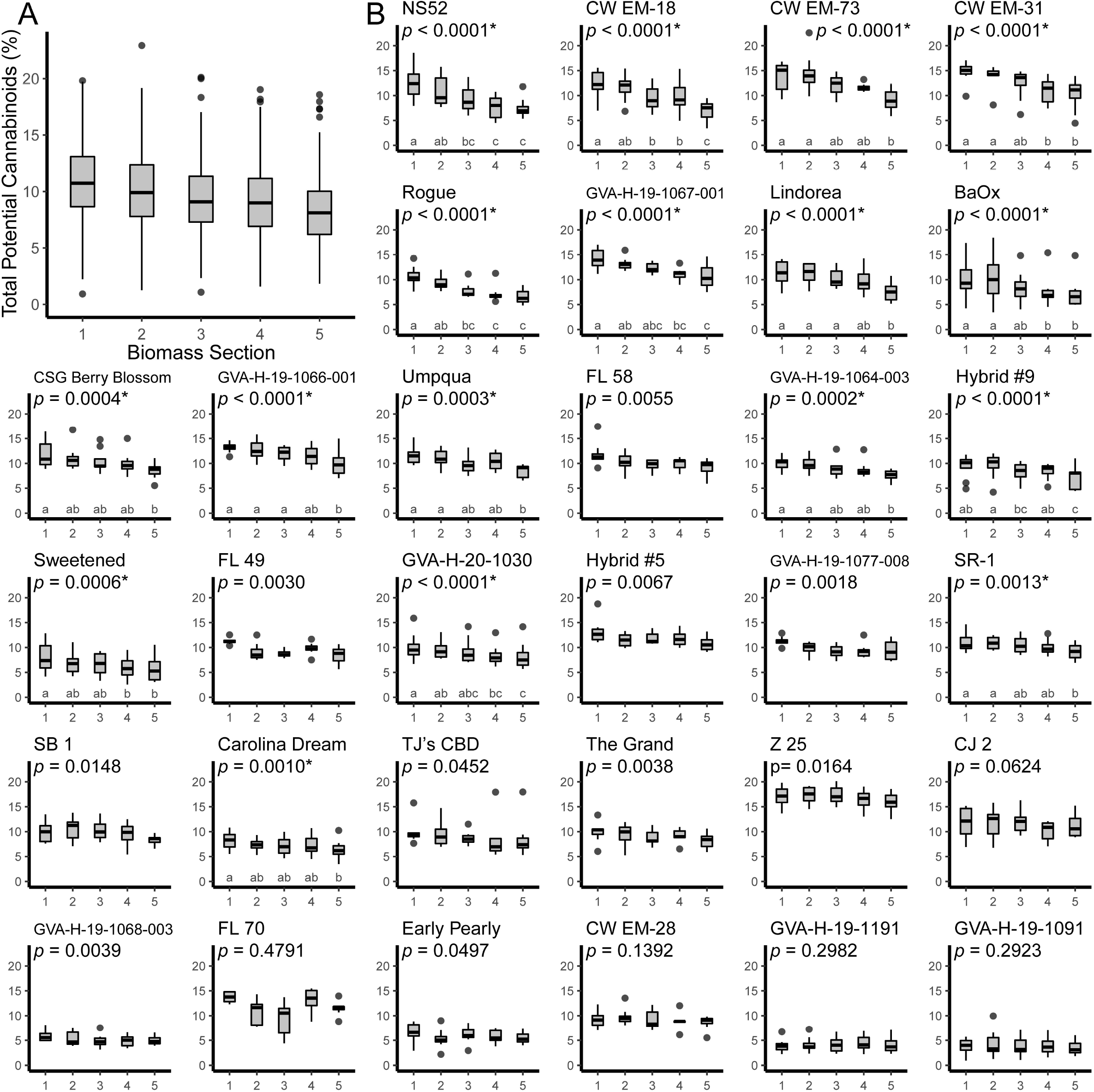
Boxplots showing the distribution of total potential cannabinoid concentration measured in post-harvest stripped biomass samples for plants partitioned into five sections where S1 represents branches derived from the top 5^th^ of the main stem, S2 those from the second to top, etc. (A) Boxplot of all samples from all cultivars. (B) Thirty-two cultivar-level boxplots ordered by the slope of the regression predicting total potential cannabinoids using biomass section. *P*-values reported are for mixed models predicting total potential cannabinoid concentration where biomass section was a fixed effect and the plot that the sample was from was a random effect. Asterisks indicate a significant effect of biomass section after correcting for multiple testing and letters indicate statistically significant differences between sections of a cultivar based on a post-hoc Tukey’s HSD test.

### 3.5 Variations in plant biomass and architecture are correlated with intra-plant variation in cannabinoid concentration

In the multiple linear regression predicting change in cannabinoid content by plant section, the overall model, after the main effect of dry canopy density was removed, was significant (*R*^2^ = 0.30, *F*(2,29) = 6,26, *p* = 0.006) (Fig. 7A). Both canopy area (β = 1.460, *p* = 0.006) and the interaction between canopy area and the log of dry canopy density (β = -0.273, *p* = 0.003) were found to be significant predictors of the change in cannabinoid content by plant section.

**Figure 7.**
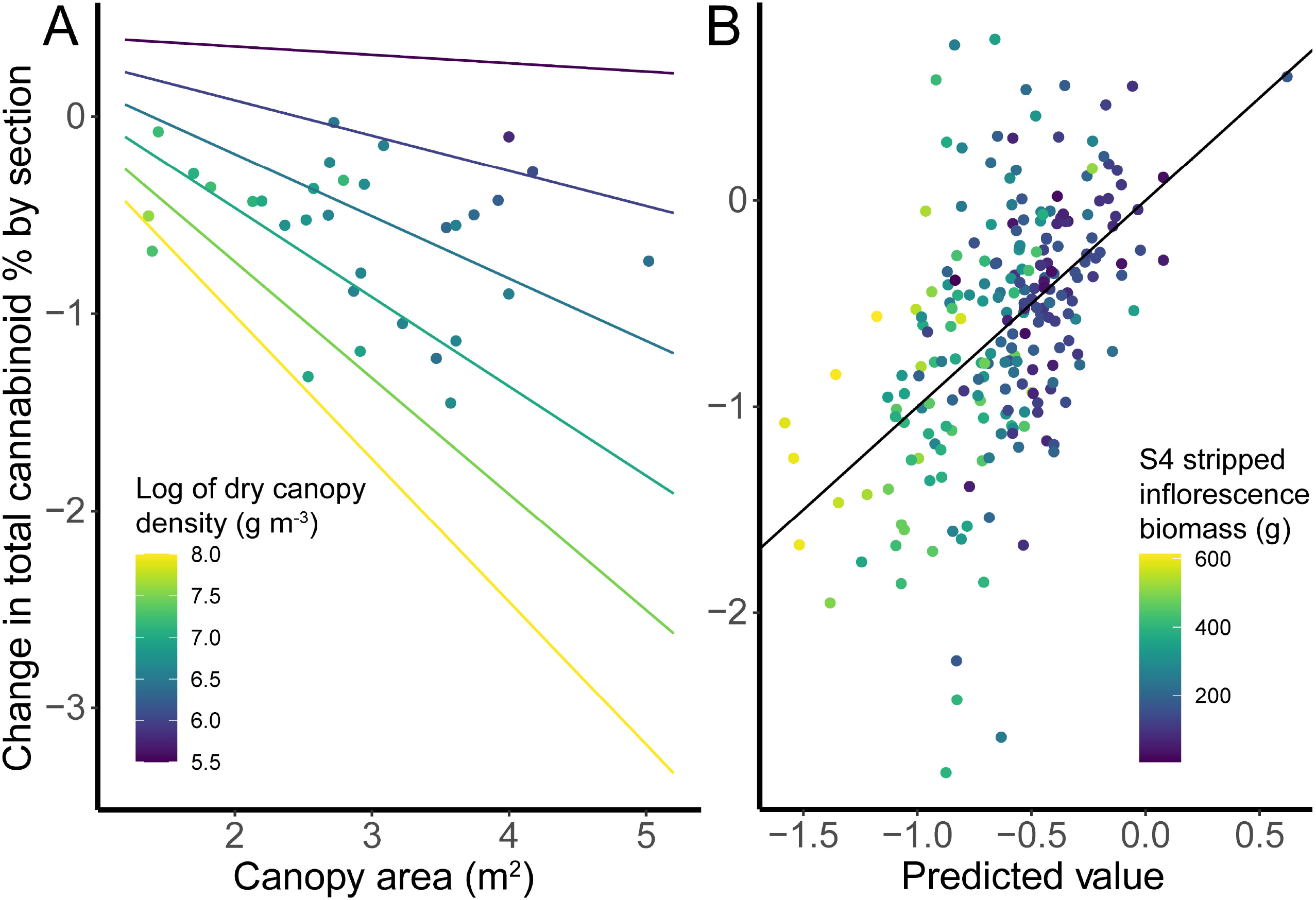
Regressions predicting changes in total cannabinoid concentration by section. (A) Multiple linear regression predicting cultivar-level values using canopy area, log of dry canopy density, and an interaction term as predictors. (B) Regression of plant-level observed values against predicted values for multiple linear regression where predictors were selected by stepwise regression. The solid black line represents a perfect correlation of predicted and observed values.

To predict the change in cannabinoid content by plant section using stepwise regression, nine predictors were identified that explained 27.7% of the variance (Fig. 7B). The regression statistics and LMG values for the nine predictors can be found in Supplementary Table S2.

Based on the LMG metric, the amount of stripped inflorescence biomass in S4 was the best single predictor, explaining 8.1% of the variance.

## 4 Discussion

### 4.1 Correlations between in-season and post-harvest cannabinoid concentrations

The strong correlation between cannabinoid concentration early in inflorescence development and concentrations in bulk stripped biomass suggests that the metabolic mechanism modulating variation in cannabinoid concentration among cultivars is initiated soon after the onset of flowering. This trend may persist earlier into floral development, with tissue accumulating greater concentrations of cannabinoids after floral induction, but before the visible onset of flowering. Richins et al. (2018) reported a significant correlation between THC concentrations in vegetative fan leaves and inflorescence material (*R*^2^ = 0.358) even before the induction of flowering.

While there is a substantial and growing body of data concerning cannabinoid accumulation after the visible onset of flowering (Pacifico *et al*., 2008; De Backer *et al*., 2012; Aizpurua-Olaizola *et al*., 2016; Richins *et al*., 2018; Yang *et al*., 2020; Stack *et al*., 2021), far less is known about variation and accumulation of cannabinoids prior to this developmental transition. With a better understanding of intra-plant cannabinoid variation prior to flowering, and the relationships between pre-inflorescence and at-harvest cannabinoid concentrations, we could broaden our mechanistic understanding of cannabinoid biosynthesis and could uncover powerful early-cycle selection criteria to breed improved cultivars. Additionally, the strong correlations between samples throughout the course of floral development could be used as a predictive tool for high-cannabinoid hemp producers to schedule regulatory testing while remaining compliant for THC concentration.

### 4.2 Distribution of inflorescence biomass within the plant canopy and relationships with plant architecture

As inflorescence biomass is the primary product of interest when producing high-cannabinoid *C. sativa*, the position and distribution of this biomass throughout the canopy is critical for growers. Based on the data presented here, growers should prefer plants that produce a greater proportion of biomass in the upper canopy. Plants that had a greater proportion of biomass in S5 produced less inflorescence biomass per unit area (Fig. 5C), and plants with a large bicone volume also had a significantly lower harvest index, potentially due to the additional stem biomass needed to support the large basal branches (Fig. 4B).

Moving toward a high-density production system, where individual plants are smaller but production per unit area is greater, has been effective in increasing yields across dozens of cropping systems (Duvick, 2005; Assefa *et al*., 2018; Postma *et al*., 2021). Through tandem improvement of cultural practices for high-density cultivation, as well as development of cultivars selected for high-density production, cultivators have been able to increase yields and production efficiency. While the bulk of this discussion focuses on outdoor production systems, similar principles govern increasing production density in controlled environment systems (Rodriguez *et al*., 2007; Amundson *et al*., 2012). Selection strategies towards improved high-cannabinoid cultivars for indoor production should consider optimization of biomass distribution for high-density cultivation systems, specifically by reducing the need for labor-intensive manual pruning and training.

### 4.3 Cultivar-dependent intra-plant variation in cannabinoid concentration

In contrast with previous studies of intra-plant variation in cannabinoid concentration (Richins *et al*., 2018; Bernstein *et al*., 2019; Danziger and Bernstein, 2021; Crispim Massuela *et al*., 2022), many of the cultivars evaluated in this study had little variance in cannabinoid concentration across the canopy. This incongruence could be attributed to many factors, including the different genotypes used in the studies, variation in sampling methodology, and differences in cultivation environment. Nevertheless, all cultivars with significant variation among sections followed the same general pattern described in the literature, with the greatest concentrations of cannabinoids accumulating in the upper-most part of the canopy.

The degree of variation in total cannabinoid concentration throughout the canopy is particularly important to growers in the context of regulatory compliance. Because THCA and CBDA are synthesized in predictable ratios by the CBDA synthase enzyme in chemotype III *C. sativa* (Zirpel *et al*., 2018; Toth *et al*., 2020, 2021; Yang *et al*., 2020; Stack *et al*., 2021), remediation of non-compliant floral biomass by blending it with plant tissue containing lower THC concentration is effective primarily due to differences in the total concentration of cannabinoids among plant sections. If there is not substantial variation in cannabinoid concentration in stripped inflorescence biomass by section, dilution with inflorescence biomass from lower sections of the canopy would not reduce the total THC concentration sufficiently to bring it below the compliance threshold.

### 4.4 Correlations with intra-plant variation in cannabinoid concentration

The factors underpinning the variation, or lack of variation, in cannabinoid concentration throughout the canopy must be understood to develop efficient production systems. This dataset complements previous work investigating the impact of plant architecture manipulation on intra-plant variation in cannabinoid accumulation (Danziger and Bernstein, 2021; Crispim Massuela *et al*., 2022). Specifically, given that manipulation of plant architecture and canopy density can cause local changes in cannabinoid concentration, does existing variation in plant architecture among cultivars correlate with intra-plant variation in cannabinoid content? Interestingly, there was a significant interaction between canopy area and dry canopy density in predicting decrease in cannabinoid concentration. Plants with a small canopy area, independent of canopy density, showed very little reduction in cannabinoid concentration, while plants with a greater canopy area exhibited canopy density-dependent decreases in cannabinoid concentration. This lends support to hypotheses that factors including light penetration and density of non-inflorescence leaves influence intra-plant variation in cannabinoid concentration. In many species, variation in light penetration has been shown to contribute to intra-plant variation in secondary metabolites, including: flavonoids (Del Valle *et al*., 2018), anthocyanins (González-Talice *et al*., 2013), and tannins (Mole *et al*., 1988). Additional work examining light penetration into the canopy, photosynthetic activity throughout the canopy, and sub-canopy lighting will provide a better understanding of the degree to which variability in light intensity modulates variation in cannabinoid concentration.

Stepwise regression modeling indicated that stripped inflorescence biomass produced in S4 was the best single predictor of intra-plant variation cannabinoid content, such that plants with greater amounts of stripped inflorescence biomass in S4 had a greater decrease in cannabinoid concentration moving down the plant. The next most important predictors were the amount of biomass in S3, the proportion of S4 that comprised stripped inflorescence biomass, and the kite angle ratio. Of the nine predictors in the final model, six were biomass measurements, while the other three were plant architecture measurements. Though it was not selected as a predictor in the final model, the relationships between the selected plant architecture and biomass measurements with canopy density suggest that variation in micro-environments throughout the plant canopy drives variation in cannabinoid concentration. Plants that produce less stripped inflorescence biomass and have a lower relative MCDH could have a more homogenous and open canopy environment (light, temperature, humidity, airflow, etc.) and thus a more homogeneous cannabinoid profile throughout the canopy.

Further studies are needed to characterize the interactions between genotype, environment, planting density, and manipulation of plant architecture, specifically contrasting genotypes with and without innate intra-plant variation in cannabinoid concentration.

### 4.5 Is intra-canopy variation in cannabinoid concentration adaptive?

Based on this study and others, intra-plant variation in cannabinoid concentration correlates with variability in photosynthate production driven by microclimates within the canopy. However, there are alternative, and not mutually exclusive, hypotheses that could explain intra-plant variation in cannabinoids. For example, assuming that cannabinoids confer protection to proximal tissues, adaptive variation in cannabinoid concentration among inflorescences from various positions within the canopy is consistent with optimal defense theory (Pavia *et al*., 2002). Cannabinoid synthesis is concentrated in tissues proximal to those producing seeds, and as such may be under strong selective pressure through influencing seed survival, dispersal, and subsequent reproductive success contributing to plant fitness. It is well documented in many angiosperms that seeds produced in different regions of the canopy can vary in size and biochemical composition (Hendrix, 1984; Raboy and Dickinson, 1987; Huber *et al*., 2016).

Further, intra-specific variation in seed size is widely associated with differences in seed dispersal and seeding establishment (Stanton, 1984; Baskin and Baskin, 2014; Visser *et al*., 2016; Genna and Pérez, 2021). Thus, variation in seed provisioning and fitness throughout the *C. sativa* canopy, potentially driven by microclimatic variation, could drive natural selection for variation in cannabinoid production among inflorescences, with greater concentrations of cannabinoids being produced in inflorescences with better-provisioned seeds. Seeds and cannabinoids are both metabolic sinks and their proximity and strength relative to photosynthetic source tissues likely limits their production. This study could not capture variation in seed traits as the plants were maintained unpollinated. Little is known about intra-plant variation in seed traits in *C. sativa* and future studies, particularly studying wild or feral germplasm, which has been under natural selection, would be valuable in the development of ecological models.Broader application of these models to other species might help to dissect the basis of adaptive intra-plant variation in secondary metabolites.

## Abbreviations

(THC): Δ^9^□tetrahydrocannabinol
(CBD): Tetrahydrocannabinolic acid (THCA) Cannabidiol
(CBDA): Cannabidiolic acid
(CBCA): Cannabichromenic acid
(CBC): Cannabichromene
(CBGA): Cannabigerolic acid
(CBG): Cannabigerol
(CBN): Cannabinol
(THCV): Tetrahydrocannabivarin
(THCVA): Tetrahydrocannabivarinic acid
(CBDV): Cannabidivarin
(CBDVA): Cannabidivarinic acid
(CBL): Cannabicyclol
(CBLA): Cannabicyclolic acid
(Δ^8^-THC): Δ^8^□tetrahydrocannabinol
(MCD): Maximum canopy diameter
(MCDH): Height at maximum canopy diameter

## 5 Supplementary Data

*Table S1*. List of traits and formulas used in the study.

*Table S2*. Analysis of variance from stepwise selection as well as relative importance (R %) of predictor variables for change in cannabinoid concentration by section (n=226).

*Fig. S1*. Flowering stage rating scale for pistillate *C. sativa* following the descriptors of Carlson et al. (2021).

*Fig. S2*. Elbow plot for *k*-means clustering of cultivars by dry biomass proportions.

*Dataset S1*. Plot-level summary statistics for all traits.

*Dataset S2*. HPLC data showing cannabinoid concentrations from 1w, 3w, and 5w regulatory shoot tip samples.

*Dataset S3*. HPLC data showing cannabinoid concentrations from biomass section subsamples.

## 6 Acknowledgements

We are grateful to the research teams from the L. Smart, C. Smart, Viands, Rose, and Bergstrom labs for their excellent technical assistance on this project, especially Deanna Gentner, Stephen Snyder, Allison DeSario, Teagan Zingg, Savanna Shelnutt, Emily McFadden, Lauren Carlson, Colin Day, Garrett Giles, Dr. Heather Grab, Nicholas Kaczmar, Jesse Chavez, Ryan Crawford, Jason Schiller, Johanna Gertin and the farm crews at Cornell AgriTech and the CUAES campus area farms. We are also grateful to Stem Holdings Agri, Sunrise Genetics, Phytonyx, Industrial Seed Innovations, Castetter Sustainability Group, Ryes Creek, Front Range Biosciences, Charlotte’s Web, Blue Forest Farms, Wessels’ Farm, and Boring Hemp for generously contributing cultivars.

## 7 Author Contributions

GMS: conceptualization, data curation, formal analysis, investigation, methodology, writing – original draft preparation. CHC, JAT: investigation, methodology. JLC: investigation. GP: methodology. JKCR, DRV, JLH, LBS: funding acquisition, supervision. All authors contributed to writing – review and editing.

## 10 Data Availability

All data supporting the findings of this study are available within the paper and within its supplementary materials published online.

